# Redox-active di-O-methylated coumarins exudation contributes to genotype-dependent iron deficiency tolerance in soybean

**DOI:** 10.64898/2026.03.26.714459

**Authors:** Francisco José Jiménez-Pastor, Edgar García-Cruz, Raúl Bouzada-Díaz, Javier Abadía, Jorge Rodríguez-Celma, Ana Álvarez-Fernández

**Affiliations:** Plant Micronutrient Stress Physiology Group, Estación Experimental de Aula Dei (EEAD), CSIC, Avenida de Montañana 1005, E-50059 Zaragoza, Spain

**Keywords:** Catechol, Coumarins, Genotypic variation, Iron deficiency chlorosis, Iron efficiency, Iron mobilization, Methylsideretin, Rhizosphere chemistry, Root exudates, Soybean

## Abstract

Iron (Fe) deficiency is a widespread disorder limiting global soybean (*Glycine max* (L.) Merr.) production. Although root exudation is a key adaptive mechanism for Fe scarcity in species like Arabidopsis, a detailed chemical characterization of soybean exudates is lacking. Here, we examined the accumulation and secretion of phenolic compounds in soybean roots and their correlation with intraspecific tolerance to Fe-deficiency chlorosis. Seven soybean genotypes with contrasting tolerance, derived from U.S. breeding programs, were analyzed. Root exudates from Fe-deficient soybean plants solubilized ferric oxide. We identified and quantified 28 coumarin-type phenolics, with catechol methylsideretin as the predominant component. Although the qualitative coumarin profile was consistent across all genotypes, Fe-efficient lines secreted these compounds at higher levels or earlier during Fe deficiency than Fe-inefficient lines. The efficient genotype A7 showed coordinated upregulation of coumarin biosynthesis and secretion, whereas this response was weaker in the Fe-inefficient genotype IsoClark. Catechol methylsideretin concentrations strongly correlated with the ability of root exudates to mobilize Fe from ferric oxide. The conserved phenolic profile, together with divergence from those reported in non-legume species, suggests lineage-specific adaptations and ecological roles beyond Fe mobilization. These results highlight genotype-dependent exudation as a determinant of soybean Fe-deficiency tolerance, with implications for breeding.

**HIGHLIGHT:** Iron deficiency induces soybean root exudates containing predominantly catechol methylsideretin which mobilize iron; genotypes differing in Fe efficiency show conserved qualitative but contrasting quantitative coumarin profiles.

## INTRODUCTION

Soybean (*Glycine max* (L.) Merr.) is the most widely cultivated legume worldwide and a major oilseed crop of high economic value. It plays a central role in global food supply, trade, and sustainable agriculture. Due to its high nutritional density, soybean provides a key source of plant-based protein (30–35%) and dietary iron (Fe) (Usman et al., 2025). Through symbiosis with rhizobia, it also contributes substantially to nitrogen (N) cycling, accounting for ∼16 of the 21 million tons of N fixed by legumes annually (Herridge et al., 2008). Despite its agronomic and nutritional importance, soybean yield is frequently limited by Fe deficiency chlorosis (IDC), a widespread disorder in major soybean-producing regions, particularly on calcareous soils, causing yield losses up to USD 260 million per year in the United States (Merry et al., 2022).

Iron is essential for plant life, participating in photosynthesis, respiration, and DNA synthesis. In legumes, its role is amplified as a cofactor in nitrogenase and leghemoglobin, both essential for symbiotic N fixation (González-Guerrero et al., 2023), and as a regulator of nodulation signaling pathways (Ren et al., 2025). Although Fe is the fourth most abundant element in the Earth’s crust, it is poorly available to plants because Fe³□ forms insoluble hydroxides in aerated, neutral to alkaline soils (Lindsay et al., 1995). Plants have evolved two phylogenetically separated strategies to mine Fe from soil. Graminaceous species, such as major cereals, use a chelation-based strategy, synthesizing and releasing into the rhizosphere mugineic acid (MA)–type phytosiderophores that solubilize ferric oxides as Fe³L–MA chelates, which are subsequently taken up by root cells (Kobayashi and Nishizawa, 2012). Rice and maize predominantly release 2’-deoxyMA, whereas Fe-efficient species such as barley exudes a more diverse array of MAs. Nongraminaceous species, such as *Arabidopsis thaliana* and soybean, mine Fe from soil via a reduction-based strategy that involves the mobilization and reduction of Fe^3+^ to Fe^2+^ prior to uptake by root cells. Molecular components associated with this strategy have been characterized in *A. thaliana*. The enzymatic reduction of ferric chelates to Fe²□ at the root surface is catalyzed by the plasma membrane (PM)-localized ferric reductase FERRIC REDUCTION OXIDASE 2 (FRO2) (Robinson et al., 1999), and the resulting Fe²□ is subsequently transported into root cells by the PM transporter IRON-REGULATED TRANSPORTER 1 (IRT1) (Vert et al., 2002). Rhizosphere acidification driven by the PM HL-ATPase AHA2 enhances both Fe³□ solubility and FRO2 activity, which is optimal at acidic pH (5.0–5.5) and becomes nearly inactive above pH 7 (Santi and Schmidt, 2009). Under alkaline and aerobic conditions, Fe³□solubility drops dramatically and Fe²□is easily reoxidized, reducing the efficiency of this strategy.

Roots of nongraminaceous species also release small molecules, originally described as “reductants” (1960s–1990s) for their ability to reduce Fe³□to Fe²□. Based on available analytical techniques, the compounds were consistent with phenolic and flavin-type metabolites (Römheld and Marschner, 1986; Rodríguez-Celma et al., 2011). Their specific molecular identities and physiological significance remained unclear for decades, until being elucidated mainly in *A. thaliana*. Under limited Fe availability, *A. thaliana* roots synthesize and secrete coumarin-type phenolics that mobilize Fe from sparingly soluble sources, supporting plant fitness (Rodríguez-Celma et al., 2013; Fourcroy et al., 2014; Schmid et al., 2014; Schmidt et al., 2014; Sisó-Terraza et al., 2016a; Rajniak et al., 2018). This class of phenolics comprises over 1,300 structures in plants (Borges et al., 2005). Those produced by Fe-deficient *A. thaliana* share the scopoletin backbone, with sideretin and fraxetin—both catechols, a chemical group capable of reducing and chelating Fe^3+^—being the most abundant (Fig. 1). External pH modulates coumarin roles: at slightly acidic pH, sideretin dominates and, together with FRO2, provides Fe^2+^ for uptake (Rajniak et al. 2018; Paffrath et al., 2024); at circumneutral pH, fraxetin dominates and mobilizes Fe from sparingly available ferric sources providing soluble Fe^3+^ complexes to the FRO2/IRT1/AHA2 system (Paffrath et al., 2024; Robe et al., 2025). In line with that, intraspecific differences in fraxetin secretion were found to correlate with *A. thaliana* adaptation to soil carbonate (Tsai et al., 2018; Terés et al., 2019), and fraxetin can enhance Fe nutrition even in *irt1* plants and in the presence of strong Fe²□chelators, suggesting uptake of Fe^3+^–fraxetin complex akin to Fe³□-MA in grasses (Robe et al., 2025).

**Fig. 1.**
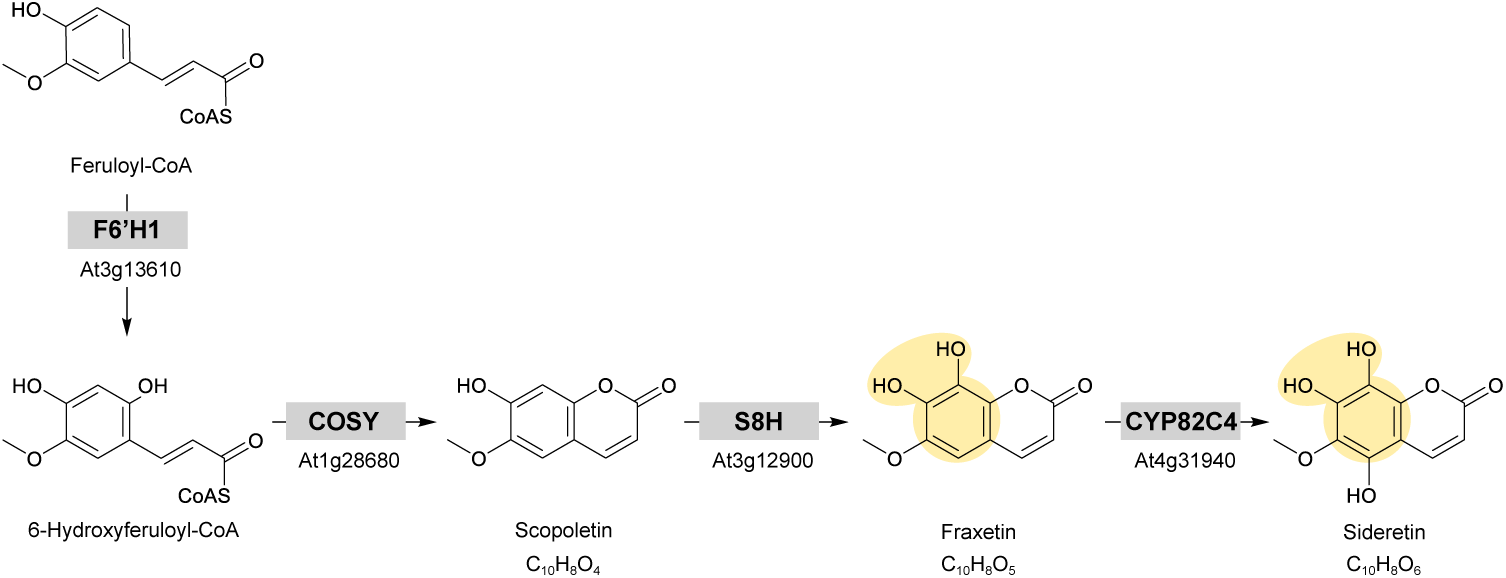
Coumarin biosynthesis pathway induced by iron deficiency in *Arabidopsis thaliana* roots. Feruloyl-CoA from the general phenylpropanoid pathway is first *ortho*-hydroxylated by Feruloyl-CoA-6’-Hydroxylase 1 (F6’H1) to form 6-hydroxyferuloyl-CoA, which is converted to the coumarin scopoletin by COumarin SYnthase (COSY). Sequential hydroxylations catalyzed by Scopoletin 8-Hydroxylase (S8H) and cytochrome P450 82C4 (CYP82C4) produce the catechol coumarins, fraxetin and sideretin. Chemical structures are shown for all metabolites; molecular formulae are indicated for coumarins only. The catechol moiety is highlighted in yellow.

These coumarins are synthesized via a pathway beginning with ortho-hydroxylation of feruloyl-CoA by the 2-oxoglutarate-dependent dioxygenase FERULOYL-CoA 6’-HYDROXYLASE 1 (F6’H1; Fig. 1) (Schmid et al., 2014; Schmidt et al., 2014). The resulting 6-hydroxyferuloyl-CoA is converted into scopoletin by the BAHD acyltransferase COUMARIN SYNTHASE (COSY) in dark-grown tissues (Vanholme et al., 2019). Scopoletin is sequentially hydroxylated to fraxetin and sideretin by SCOPOLETIN 8-HYDROXYLASE (S8H) (Tsai et al., 2018) and cytochrome P450 enzyme CYP82C4 (Rajniak et al., 2018), respectively. Fraxetin and sideretin secretion into the rhizosphere is mediated by the PLEIOTROPIC DRUG RESISTANCE 9/ATP-BINDING CASSETTE G37 PDR9/ABCG37 transporter (Fourcroy et al., 2014; Ziegler et al., 2017). Expression of F6’H1, S8H, CYP82C4, along with IRT1 and FRO2, is regulated by the bHLH transcription factor FER-LIKE IRON DEFICIENCY-INDUCED TRANSCRIPTION FACTOR (FIT) and strongly induced under Fe deficiency (Schmid et al., 2014; Rajniak et al., 2018). Orthologs of these genes exist across many nongraminaceous species, indicating a broadly conserved biosynthetic module (Rajniak et al., 2018). Coumarin secretion has been documented in several dicots, including *Brassica rapa*, tobacco, and tomato (Rajniak et al., 2018; Lefèvre et al., 2018; Astolfi et al., 2020). Among legumes, only Medicago species have been characterized, which exude flavins but not coumarins (Rodríguez-Celma et al., 2011; Rajniak et al., 2018; Gheshlaghi et al., 2021), suggesting lineage-specific differences in the metabolites used by plants to mine Fe.

Responses to Fe deficiency in soybean include rhizosphere acidification, ferric chelate reduction, enhanced Fe uptake, and secretion of “reductants”. Acidification capacity is limited and inconsistently linked to Fe efficiency (Jolley et al., 1986; Zocchi et al., 2007), whereas ferric chelate reduction and “reductants” secretion have been consistently associated with Fe efficiency (Brown et al., 1961; Brown and Ambler, 1973; Jolley et al., 1986; Waters et al., 2018). A screening of 50 genotypes using ^14^CO_2_ showed increased total carbon exudation under Fe deficiency, with the highest among them being the Fe-efficient and responsive lines (Raj et al., 2019). In addition, genes involved in coumarin biosynthesis (*F6’H1*, *S8H* and *CYP82C4*) were upregulated in roots, with *F6’H1* expression higher in the tolerant line (Moran-Lauter et al., 2014; Waters et al., 2018). Notably, the soybean homolog of *AtF6’H1* co-localizes with a major quantitative trait locus on chromosome 3 associated with IDC tolerance, but the secreted compounds have not yet been characterized.

In this study, we characterized phenolic accumulation and secretion in soybean roots under Fe deficiency and their association with intraspecific IDC tolerance. We identified 28 coumarins, with catechol methylsideretin as the predominant compound—a pattern distinct from other legumes, which secrete flavins, and from non-legumes such as *A. thaliana* and tobacco. Iron-efficient lines secreted higher amounts of coumarins and/or initiated secretion earlier than inefficient lines. The Fe-mobilizing capacity of root exudates, especially at circumneutral pH, strongly correlated with catechol methylsideretin abundance. These results indicate that catechol methylsideretin secretion is a root trait specific to soybean and a promising target to improve tolerance to IDC.

## MATERIALS AND METHODS

### Plant materials and growth conditions

Soybean (*Glycine max* (L.) Merr.) genotypes with contrasting Fe deficiency tolerance were used: three Fe-inefficient (IsoClark, Anoka and B216) (and five as Fe-efficient (Clark, A15, A97, AR3 and A7) (Jolley et al., 1986; Bernard et al., 1991; Peiffer et al., 2012). Several of these genotypes are closely related: A7 and A15 were derived monoparentally from AP9; A97 (*Northup Kin S20-20 x Asgrow A2234*) is a parent of AR3 (*A97 x Pioneer P9254*); Clark (*Lincoln(2) x Richland*) is a parent of IsoClark (*Clark x T203*). In contrast, B216 (*Corsoy x Wayne*) is not directly related to any of the other genotypes. Seeds were kindly provided by Prof. Silvia Cianzio (Iowa State University, U.S.). They were germinated, pre-grown and grown in a controlled-environment walk-in chamber (Fitoclima 10000 EHHF, Aralab, Albarraque, Portugal) under a 16/8 h day/night regime, 25 °C, 70% relative humidity, and a photosynthetic photon flux density of 350 □mol m^-2^ s^-2^. Growth and treatment conditions were optimized based on preliminary experiments as described in Note S1. For the results shown here, seeds were germinated and pre-grown in vermiculite for 8 d. Seedlings (with cotyledons fully expanded at this stage) were then grown for 3 d, in 10 L plastic bucket (20 seedlings per bucket) with a continuously aerated ½ Hoagland nutrient solution pH 5.5, with 25 µM Fe^3+^EDDHA. Then, cotyledons were removed, and seedlings were grown for 13 d more in 0.5 L plastic pot (1 seedling per pot) filled with a continuously aerated ½ Hoagland nutrient solution pH 5.5, with 0 (−Fe) or 25 µM Fe^3+^EDDHA (+Fe). The pH was daily adjusted to 5.5 with NaOH or HNO_3_ as required. Three independent batches were grown and analyzed for IsoClark, Clark, A15, and A7, and a single batch for B216, A97, and AR3. Replicate numbers ranged from 4–14 in −Fe and 3–6 in +Fe treatments, depending on genotype. Pots without plants, containing aerated nutrient solution (with or without Fe), were also placed in the growth chamber and their nutrient solutions sampled and used as in pots with plants.

During treatments, nutrient solution pH and leaf SPAD values (young and old leaves) were recorded daily. Aliquots of nutrient solution (8 mL) were collected daily, filtered (0.45 µm PVDF), and stored at –20 °C or –80 °C. Whole roots were harvested at various time points, weighed, flash-frozen, and stored at –80 °C for phenolic/flavin extraction. Shoot and root fresh mass was recorded for each plant. Root FCR activity was measured daily in IsoClark and A7. In a subset of A7 plants (+Fe and –Fe), organs were also collected for mineral analysis (Sisó-Terraza et al., 2016b).

### Iron mobilization from a ferric oxide promoted by root exudates

The ability of nutrient solutions from −Fe plants to mobilize Fe from a ferric oxide was assessed following Sisó-Terraza et al. (2016a, b). For each genotype, equal volumes of nutrient solution collected from –Fe plants between days 5 and 13 were pooled, vacuum-dried, and reconstituted in incubation buffer to obtain a 20× concentrate. Assays contained 10 mg of poorly crystalline ferric oxide and 1.5 mL of reaction medium consisting of 4× concentrated pooled exudates, 0 or 1 mM NADH, 0 or 600 µM BPDS, and 25 mM ammonium acetate (pH 5.5) or 25 mM HEPES-KOH (pH 7.5). Assay blanks and nutrient-solution blanks from pots without plants were included. Mixtures were incubated in the dark at 25□°C for 6□h under constant agitation (300 rpm) in a Thermomixer Comfort (Eppendorf, Hamburg, Germany). After incubation, solutions were filtered (0.22 µm PVDF) and Fe²□–BPDS□ was quantified by absorbance at 535 nm (ε = 22.14 mM□¹ cm□¹). Total Fe was measured by ICP-MS after 1:1 dilution in 6% HNOL (TraceSELECT Ultra, Sigma–Aldrich). All assays were performed in duplicate or triplicate and results are reported as mean ± SD.

### Extraction and analyses of phenolic compounds from roots and nutrient solutions

Phenolic compounds were extracted from roots following the method of Sisó-Terraza *et al*. (2016a), with minor modifications. Frozen roots were ground to a fine powder in liquid N_2_ using a mortar and pestle. Approximately 100 mg aliquots were further pulverized using a Retsch M301 ball mill (Restch, Düsseldorf, Germany) for 3 min and subsequently homogenized three times with 1 mL of 100% LC-MS grade methanol (LiChrosolv®, Merck) in the same mill for 5 min each. The three resulting supernatants were pooled and dried using a SpeedVac device (MiVac, GeneVac^TM^, Ipswich, UK). The dried residue was immediately reconstituted in 250 µL of the initial mobile phase (5% v/v methanol and 0.1% v/v formic acid). For nutrient solutions, 8 mL aliquots were vacuum-dried, and the resulting residue was dissolved in 400 µL of the initial mobile phase. Prior to analysis, all root extracts and concentrated nutrient solutions were filtered through 0.22 µm PVDF ultrafree-MC centrifugal filter devices (Merck-Millipore).

An untargeted analysis was performed using high-performance liquid chromatography coupled to ultraviolet–visible spectrophotometry and electrospray ionization time-of-flight mass spectrometry [HPLC-UV/VIS-ESI-MS(TOF)]. The system consisted of an Alliance 2795 HPLC (Waters, Milford, MA, USA) coupled to a PDA 2996 UV/VIS (Waters) detector and a MicrOTOF mass spectrometer (Bruker Daltonics, Bremen, Germany) equipped with an electrospray (ESI) source. Chromatographic separations were carried out following the protocol of Sisó-Terraza et al. (2016a), with slight modifications to the original elution program (details in Supplementary Table S1). The ESI-MS(TOF) settings were as described by Sisó-Terraza et al. (2016a). The system was operated using microTOF control 5.5.9.8 and HyStar 3.2 SR4 (Bruker Daltonics).

For compound annotation, selected samples were also analyzed using the Alliance 2795 HPLC coupled to an ion trap mass spectrometer (HCT Ultra, Bruker Daltonics) equipped with an ESI source. The same HPLC conditions described above were used, and ion trap operating settings are provided in Supplementary Table S2. Ions of interest were subjected to collision-induced dissociation (CID) using hellium as the collision gas for 40 ms, to generated successive sets of fragment ion sets (MS^n^) (Sisó-Terraza et al., 2016a). Instrument control was performed using Esquire 1.2.214.16 and HyStar 3.2 SR4 software (Bruker Daltonics). Data were processed with Data Analysis 4.2 SR1 (Bruker Daltonics). Molecular formulae were assigned using the Smart Formula tool in Data Analysis, based on < 5 ppm mass accuracy and SigmaFit^TM^ criteria.

For quantification, root extracts and concentrated nutrient solutions were supplemented with internal standards (ISs) and 5 mM NADH immediately before analysis using the same HPLC-UV/VIS-ESI-MS(TOF) system and operating conditions. Potential co-elution of isobaric compounds with ISs was assessed beforehand to avoid interferences. The following compounds were used as ISs: artemicapin C (a methylenedioxy-coumarin) for the quantification of coumarin aglycones; esculin (the glucoside form of the coumarin esculetin) for the quantification of coumarin hexosides; isoferulic acid (a positional isomer of ferulic acid) for the quantification of ferulic acid; and caffeic acid for the quantification of non-canonical hydroxycinnamic acids (HCAs), 3-(trihydroxymethoxyphenyl)acrylic acid, and 3- (dihydroxydimethoxyphenyl)dioxopropanoic acid. ISs concentrations in the injected solutions were: 20 µM artemicapin C, 20 µM esculin, 15 µM isoferulic acid and 40 µM caffeic acid. Analyte concentrations were determined using external calibration with internal standardization. Peak areas for analytes, standards and ISs were obtained from extracted ion chromatograms corresponding to the [M+H]^+^ *m/z* ±0.05. Exceptions included glycosides, for which [M-hexose+H]^+^ ions were used, and hydroxycinnamic acids, for which [M-H_2_O+H]^+^ ions were used. Scopolin, fraxin (and isomers), esculetin, fraxetin (and isomers), scopoletin, isofraxidin, ferulic acid, and fraxinol were quantified using authenticated standards. Coumarins lacking available commercial standards were quantified as fraxetin (aglycones) or fraxin (hexosides). Concentrations of the non-canonical HCAs described above were estimated as 5-hydroxyferulic acid equivalents.

### Gene expression analysis

Genes analyzed here were previously reported as Fe deficiency-responsive in soybean by Peiffer et al. (2012), Moran-Lauter et al. (2014), and Waters et al. (2018). Root tissues from A7 and IsoClark were sampled at 3, 4, and 5 days after treatment onset. Three biological replicates were used for the −Fe treatment and two for the +Fe treatment. Total RNA was extracted with Tri-Reagent (Life Technologies, Carlsbad, CA, USA) and treated with Turbo^TM^ DNase (Life Technologies). RNA integrity was confirmed using agarose gel electrophoresis. cDNA was synthesized from 1 µg of DNase-treated RNA using the Transcriptor First Strand cDNA synthesis kit (Roche Diagnostics, Mannheim, Germany). Quantitative real-time RT-PCR (RT-qPCR) was performed on a 7500 Real-Time PCR System (Applied Biosystems, Foster City, CA, USA) with Power SyBR^TM^ Green Master Mix (Applied Biosystems). Primers are listed in Supplementary Table S3. Transcript levels were normalized to the *G. max peptidyl-prolyl cis-trans isomerase* gene (CYP2; *GLYMA_12G024700*) and calculated as Normalized Relative Quantity (NRQ) following Hellemans et al. (2007).

### *In vivo* root ferric chelate reductase activity

Ferric chelate reductase (FCR) activity was measured in freshly excised root sections (∼70 mg; 0–35 mm from the root tip) by monitoring Fe^2+^(BPDS)_3_ formation from Fe^3+^-EDTA (Bienfait et al., 1983). Sections, sampled 2–3 h after light onset, were incubated for 60 min at room temperature in the dark in 1 mL of assay solution containing 500 μM Fe^3+^-EDTA and 300 μM BPDS, buffered at pH 5.5 with 10 mM MES. To estimate the contribution of soluble reductants released from roots, sections were incubated in assay medium without Fe^3+^-EDTA; then, sections were removed and Fe^3+^-EDTA added. Aliquots were filtered through 0.45 µm PVDF ultrafree-MC centrifugal filters (Millipore) at 10,000 g for 3 min. Absorbance was measured at 535 nm [to quantify Fe^2+^-(BPDS)_3_] and 735 nm (to correct for turbidity) using a Shimadzu UV-2450 (Tokyo, Japan). Fe^2+^-(BPDS)_3_ concentrations were calculated using a molar extinction coefficient of 22.14 mM^−1^ cm^−1^.

### Mineral analysis

Dried plant tissue (∼200 mg) was digested with ultrapure 19% HNO_3_ and 8% H_2_O_2_ for 55 min at 190° C in an Ethos 1 microwave oven (Milestone, Sorisole, Italy). Fe, Mn, Cu and Zn concentrations were determined by flame atomic spectrometry using a SOLAAR 969 instrument (Thermo, Cambridge, UK).

### Statistical analyses

All analyses were performed in R (v4.3.1; R Foundation, Vienna, Austria). Phenolic compound data (nmol per plant) from roots and nutrient solutions across Fe treatments, sampling times, and seven genotypes were square root-transformed, mean-centered, and left unscaled for principal component analysis (PCA) using the *prcomp*() function. PCA scores and loadings were visualized with ggplot2. Hierarchical clustering heatmaps were generated with *pheatmap*() using Manhattan distance and average linkage. Data normality and homogeneity of variance were assessed with Shapiro–Wilk and Levene’s tests. Pairwise comparisons employed Student’s t-test or Wilcoxon–Mann–Whitney test (p < 0.05). For multiple comparisons, Tukey’s HSD was applied when variances were equal, and Games–Howell test when unequal.

## RESULTS

### Root exudates from iron-deficient soybean plants enable ferric oxide solubilization

To determine whether soybean plants release metabolites capable of mobilizing Fe from recalcitrant soil pools, seven soybean genotypes—two Fe-inefficient and five Fe-efficient—were grown hydroponically under Fe deficiency conditions for 13 days. Nutrient solutions collected from Days 5-13 were pooled by genotype and tested for their Fe-mobilizing capacity using an in vitro incubation assay with poorly crystalline ferric oxide at pH 5.5 and 7.5. Nutrient solutions from pots without plants served as blanks.

Both blanks and plant nutrient solutions showed minimal Fe mobilization on their own (<1 nmol Fe·g^-1^·min^-1^) across genotypes and pH conditions (Supplementary Fig. S1A,C). Because aeration of the hydroponic medium could oxidize root-released reductants, assays were repeated in the presence of the electron donor NADH, but results remained similar (Supplementary Fig. S1A,C). We next assessed the effect of the Fe^2+^-trapping agent BPDS, which stabilizes newly formed Fe^2+^ by forming soluble complex. Incubations containing only BPDS showed small increases in Fe mobilization (up to 3.5 nmol Fe·g□¹·min□¹; Supplementary Fig. S1C). In contrast, combining NADH and BPDS markedly enhanced Fe mobilization, reaching 1.5-15 nmol Fe·g^-1^·min^-1^ at pH 7.5 and 12-19.3 nmol Fe·g^-1^·min^-1^ at pH 5.5 (Supplementary Fig. S1B). Under these conditions, all plant nutrient solutions mobilized significantly more Fe than the blanks at pH 7.5, and genotypes B216, AR3 and A7 also did so at pH 5.5 (Supplementary Fig. S1B). BPDS captured on average 94% and 75% of the mobilized Fe at pH 5.5 and 7.5, respectively (Supplementary Fig. S1B). To better visualize genotype-differences, blank values were subtracted from plant nutrient solution data (Fig. 2). The Fe-efficient genotype A7 showed the highest mobilization rates at pH 7.5, whereas AR3 and A7 exhibited the highest rates at pH 5.5. The Fe-inefficient genotype Isoclark consistently showed the lowest mobilization.

**Fig. 2.**
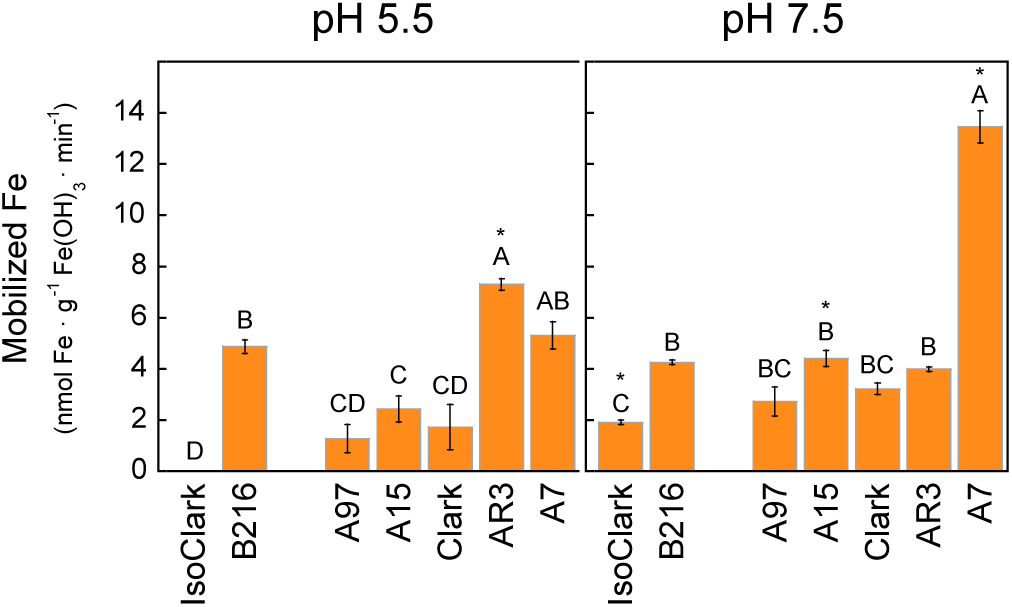
Iron mobilization from ferric oxide by root exudates from seven soybean genotypes grown under iron deficiency. Two Fe-inefficient genotypes (IsoClark and B216) and five Fe-efficient (A97, A15, Clark, AR3, and A7) were evaluated. Plants were hydroponically pre-grown for 3 d with 25□µM Fe^3+^EDDHA at pH 5.5, and then grown for 13 d in a nutrient solution without Fe, with the pH readjusted daily to 5.5. Nutrient solutions from Days 5–13 were pooled by genotype, and concentrated 20-fold. Incubation assays with 10 mg of ferric oxide for 6 h were run at two different pH values, 5.5 and 7.5, in the presence of 4-fold concentrated nutrient solution, l mM NADH and 600 µM BPDS. Total Fe mobilized was determined by ICP-MS (mean ± SE, n = 3) and corrected by subtracting Fe mobilized in blanks (nutrient solutions from pots without plants). Significant differences among genotypes at each pH (p < 0.05) are indicated by different letters above the columns. When the values obtained for the same genotype differ significantly between the two pH conditions, an asterisk marks the bar with the higher value.

These results prompted a deeper investigation into Fe deficiency-induced changes in the metabolite profiles of both roots and nutrient solutions across soybean genotypes.

### Iron deficiency induces the accumulation and release of oxygenated coumarins in soybean roots

Metabolite profiles of nutrient solutions and root extracts of Fe-sufficient (+Fe) and Fe-deficient (−Fe) plants were analyzed at multiple time points using untargeted HPLC-UV/VIS-ESI-MS(TOF). Initial data exploration focused on 320 nm absorbance chromatograms. Although all samples were examined, Fig. 3 shows representative chromatograms from Day 5 of genotype A7, selected for its high Fe-mobilizing capacity at pH 7.5 (Fig. 2). Each chromatographic peak may correspond to one or more metabolites (see Supplementary Note S2 for identification details). Chromatograms of +Fe and –Fe blanks were flat, showing no detectable peaks. In contrast, nutrient solutions from –Fe plants displayed several new peaks and strong increases in others that were minor under +Fe conditions. Root extract chromatograms were overall more complex, showing new peaks induced by Fe deficiency as well as changes in intensity of peaks present under +Fe conditions. These observations indicated that Fe deficiency alters the metabolite profiles of roots and nutrient solutions, likely involving phenolic compounds.

**Fig. 3.**
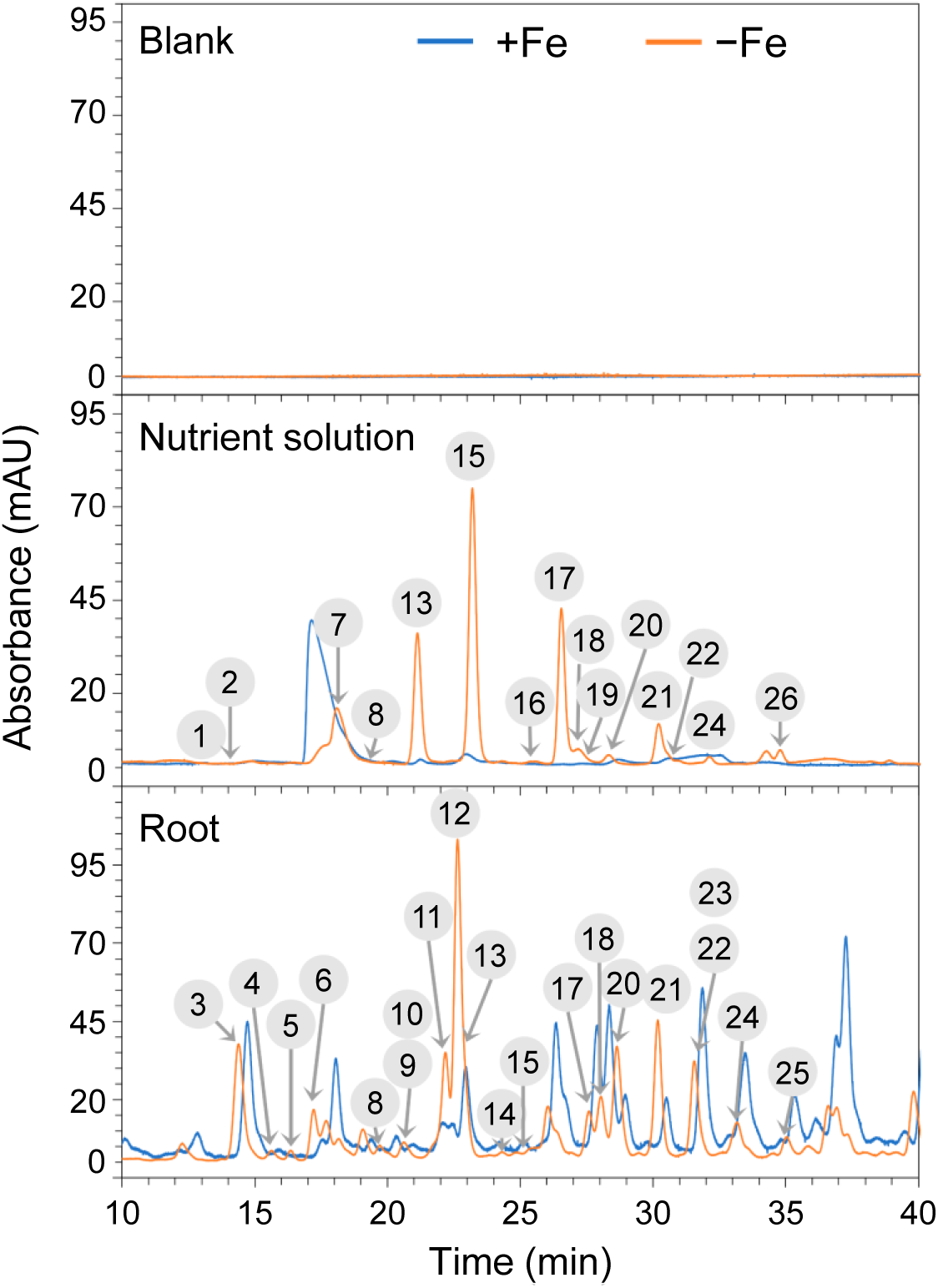
Chromatographic separation of root-accumulated and -released compounds in the A7 soybean genotype under iron-deficient and iron-sufficient conditions. Plants were hydroponically pre-grown for 3□d with 25□µM Fe³□-EDDHA at pH 5.5, and then grown for 13□d in a nutrient solution without (−Fe) or with (+Fe) 25□µM Fe³□-EDDHA, with the pH readjusted daily to 5.5. Representative 320□nm absorbance chromatograms are shown for blanks (nutrient solution from pots without plants), nutrient solutions, and root extracts from −Fe and +Fe A7 plants sampled on Day 5. Only the 10–40□min retention time window is displayed, corresponding to the annotated compounds. Encircled numbers above peaks refer to phenolic compounds listed in Table 1. mAU, milli-absorbance units.

**Table 1.** Phenolic compounds accumulated in roots and released by soybean under iron deficiency. Retention times (RT), exact mass-to-charge ratios (*m/z),* molecular formulas, and *m/z* errors (ppm) are shown. The *m/z* values correspond to [M+H]^+^ and [M-H]^-^ ions obtained from HPLC-ESI-MS(TOF) in positive and negative ionizations modes, respectively. For compounds *4* and *15* in positive mode, the *m/z* values correspond to the in-source fragment ion [M−hexose+H]^+^, which was more intense than the [M+H]^+^ ions. Grey-shaded rows indicate glycosylated compounds. Positions of the substitutions in the compound have not been confirmed by NMR studies. Common names and short names used in this study are indicated in square brackets.

| # | RT<br>(min) | Measured<br>$m/z$ | Molecular<br>formula | Calculate<br>d $m/z$ | Error<br>$m/z$<br>(ppm) | Annotation |
| --- | --- | --- | --- | --- | --- | --- |
| 1 | 12.8 | 227.0547 | $C_{10}H_{11}O_6^+$ | 227.0550 | 1.32 | 3-(trihydroxymethoxyphenyl)acrylic acid |
| | | 225.0394 | $C_{10}H_9O_6^-$ | 225.0404 | 4.44 | [dihydroxyferulic acid; sideretin degradation product] |
| 2 | 14.0 | 271.0440 | $C_{11}H_{11}O_8^+$ | 271.0448 | 2.95 | 3-(dihydroxydimethoxyphenyl)dioxopropanoic acid |
| | | 269.0308 | $C_{11}H_9O_8^-$ | 269.0303 | -1.86 | $[\alpha,\beta\text{-dioxo-2-hydroxy-5-methoxydihydroferulic acid; methylsideretin degradation product}]$ |
| 3 | 14.8 | 371.0962 | $C_{16}H_{19}O_{10}^+$ | 371.0973 | 2.96 | Dihydroxymethoxycoumarin hexoside isomer 1 |
| | | 369.0835 | $C_{16}H_{17}O_{10}^-$ | 369.0827 | -2.17 | [fraxin isomer 1] |
| 4 | 15.6 | 387.0937 | $C_{16}H_{19}O_{11}^+$ | 387.0922 | -3.88 | Trihydroxymethoxycoumarin dihexoside |
| | | 547.1278 | $C_{22}H_{27}O_{16}^-$ | 547.1304 | 4.70 | [sideretin dihexoside] |
| 5 | 16.8 | 371.0963 | $C_{16}H_{19}O_{10}^+$ | 371.0973 | 2.70 | Dihydroxymethoxycoumarin hexoside isomer 2 |
| | | 369.0840 | $C_{16}H_{17}O_{10}^-$ | 369.0827 | -3.52 | [fraxin isomer 2] |
| 6 | 18.0 | 355.1009 | $C_{16}H_{19}O_9^+$ | 355.1024 | 4.22 | 7-hydroxy-6-methoxycoumarin 7-hexoside<br>[scopolin] |
| 7 | 18.9 | 223.0240 | $C_{10}H_7O_6^+$ | 223.0237 | -1.35 | 5,7,8-trihydroxy-6-methoxycoumarin oxidized form |
| | | 221.0085 | $C_{10}H_5O_6^-$ | 221.0092 | 3.17 | [sideretin oxidized form] |
| 8 | 20.1 | 179.0334 | $C_9H_7O_4^+$ | 179.0339 | 2.79 | 6,7-dihydroxycoumarin |
| | | 177.0192 | $C_9H_5O_4^-$ | 177.0193 | 0.56 | [esculetin] |
| 9 | 20.2 | 371.0976 | $C_{16}H_{19}O_{10}^+$ | 371.0973 | -0.81 | Dihydroxymethoxycoumarin hexoside isomer 3 |
| | | 369.0824 | $C_{16}H_{17}O_{10}^-$ | 369.0827 | 0.81 | [fraxin isomer 3] |
| 10 | 20.2 | 401.1068 | $C_{17}H_{21}O_{11}^+$ | 401.1078 | 2.49 | Dihydroxydimethoxycoumarin hexoside isomer 1 |
| | | 399.0937 | $C_{17}H_{19}O_{11}^-$ | 399.0933 | -1.00 | [methylsideretin hexoside isomer 1] |
| 11 | 20.6 | 401.1069 | $C_{17}H_{21}O_{11}^+$ | 401.1078 | 2.24 | Dihydroxydimethoxycoumarin hexoside isomer 2 |
| | | 399.0937 | $C_{17}H_{19}O_{11}^-$ | 399.0933 | -1.00 | [methylsideretin hexoside isomer 2] |
| 12 | 21.0 | 209.0444 | $C_{10}H_9O_5^+$ | 209.0444 | 0.00 | 7,8-dihydroxy-6-methoxycoumarin 8-hexoside |
| | | 369.0813 | $C_{16}H_{17}O_{10}^-$ | 369.0827 | 3.79 | [fraxin] |
| 13 | 21.5 | 225.0391 | $C_{10}H_9O_6^+$ | 225.0394 | 1.33 | 5,7,8-trihydroxy-6-methoxycoumarin |
| | | 223.0238 | $C_{10}H_7O_6^-$ | 223.0248 | 4.48 | [sideretin; sideretin reduced form] |
| 14 | 22.9 | 385.1122 | $C_{17}H_{21}O_{10}^+$ | 385.1130 | 2.08 | Hydroxydimethoxycoumarin hexoside |
| | | 223.0594 | $C_{11}H_{11}O_5^+$ | 223.0601 | 3.13 | [methylfraxetin hexoside] |
| 15 | 24.6 | 209.0440 | $C_{10}H_9O_5^+$ | 209.0444 | 2.39 | 7,8-dihydroxy-6-methoxycoumarin |
| | | 207.0295 | $C_{10}H_7O_5^-$ | 207.0299 | 1.93 | [fraxetin] |
| 16 | 25.1 | 237.0389 | $C_{11}H_9O_6^+$ | 237.0394 | 2.11 | 7,8-dihydroxy-5,6-dimethoxycoumarin isomer 1 (oxidized form) |
|  |  |  |  |  |  | [catechol methylsideretin oxidized form] |
| 17 | 28.0 | 239.0542 | $C_{11}H_{11}O_6^+$ | 239.0550 | 3.35 | 7, 8-dihydroxy-5,6-dimethoxycoumarin isomer 1 |
| | | 237.0400 | $C_{11}H_9O_6^-$ | 237.0404 | 1.69 | [methylsideretin isomer 1; catechol methylsideretin] |
| 18 | 28.5 | 193.0488 | $C_{10}H_9O_4^+$ | 193.0495 | 3.63 | 7-hydroxy-6-methoxycoumarin |
| | | 191.0345 | $C_{10}H_7O_4^-$ | 191.0350 | 2.62 | [scopoletin] |
| 19 | 29.4 | 209.0437 | $C_{10}H_9O_5^+$ | 209.0445 | 3.83 | Dihydroxymethoxycoumarin isomer 1 |
| | | 207.0290 | $C_{10}H_7O_5^-$ | 207.0299 | 4.35 | [fraxetin isomer 1; isofraxetin] |
| 20 | 29.7 | 239.0543 | $C_{11}H_{11}O_6^+$ | 239.0550 | 2.93 | Dihydroxydimethoxycoumarin isomer 2 |
| | | 237.0397 | $C_{11}H_9O_6^-$ | 237.0404 | 2.95 | [methylsideretin isomer 2; methylsideretin] |
| 21 | 29.8 | 223.0590 | $C_{11}H_{11}O_5^+$ | 223.0601 | 4.93 | 7-hydroxy-6,8-dimethoxycoumarin |
|  |  |  |  |  |  | [isofraxidin; methylfraxetin isomer 1] |
| 22 | 30.7 | 223.0601 | $C_{11}H_{11}O_5^+$ | 223.0601 | 0.00 | Hydroxydimethoxycoumarin isomer 1 |
|  |  |  |  |  |  | [methylfraxetin isomer 2] |
| 23 | 31.4 | 195.0651 | $C_{10}H_{11}O_4^+$ | 195.0652 | 0.51 | 3-(4-hydroxy-3-methoxyphenyl)acrylic acid |
| | | 193.0504 | $C_{10}H_9O_4^-$ | 193.0495 | -4.66 | [ferulic acid] |
| 24 | 32.3 | 223.0600 | $C_{11}H_{11}O_5^+$ | 223.0601 | 0.45 | 6-hydroxy 5,7dimethoxycoumarin |
|  |  |  |  |  |  | [fraxinol; methylfraxetin isomer 3] |
| 25 | 33.4 | 223.0600 | $C_{11}H_{11}O_5^+$ | 223.0601 | 0.45 | Hydroxydimethoxycoumarin isomer 2 |
|  |  |  |  |  |  | [methylfraxetin isomer 4] |
| 26 | 36.3 | 223.0596 | $C_{11}H_{11}O_5^+$ | 223.0601 | 2.24 | Hydroxydimethoxycoumarin isomer 3 |
|  |  |  |  |  |  | [methylfraxetin isomer 5] |

Analyses of HPLC-ESI-MS(TOF) data focused on retention times (RTs) corresponding to Fe deficiency–dependent increases observed in 320 nm chromatograms, leading to the detection of 26 responsive ions. Accurate elemental formulas were assigned to the [M+H]^+^ and [M–H]^−^ ions or fragments (Table 1, Fig. 3), none of which were present in blanks. To obtain structural information, selected samples enriched in these compounds were analyzed by HPLC-ESI-MS(ion trap), enabling collision-induced fragmentation, and compared with authenticated phenolic standards (Supplementary Tables S4, S5). Using RT tolerance (±0.2 min), mass accuracy (≤5 ppm), and fragment pattern comparison, all 26 compounds were annotated as phenolics, predominantly coumarins (see Supplementary Note S2 for details).

Twenty-two of the 26 compounds correspond to phenolics previously reported as Fe deficiency–responsive in roots of other species (Supplementary Table S6). Among these, eight (*6*, *8*, *12*, *15*, *18*, *21*, *23*, and *24* in Fig. 3 and Table 1) matched the RTs, *m/z* values, and MSⁿ fragmentation patterns of authentic standards: scopolin, esculetin, fraxin, fraxetin, scopoletin, isofraxidin, ferulic acid, and fraxinol. Seven additional compounds (*3*, *5*, *9*, *19*, *22*, *25*, and *26*) matched the *m/z* values and fragmentation patterns—but not the RTs—of fraxin, fraxetin, and methoxyscopoletin isomers, and were annotated as positional isomers. Five metabolites (*7*, *10*, *11*, *13*, and *20*) corresponded to coumarins previously reported in Arabidopsis (Rajniak et al., 2018) and tobacco (Lefèvre et al., 2018) under Fe deficiency, including oxidized sideretin, sideretin, methylsideretin hexosides, and methylsideretin. Two metabolites (*4* and *14*) were annotated as glycosides of trihydroxymethoxycoumarin and methylfraxetin, supported by MSⁿ spectra showing neutral losses of one or two hexosyl residues. Four compounds (*1*, *2*, *16, 17*) are newly reported here: compound *17* is catechol methylsideretin (an isomer of *20*), compound *16* is the fully oxidized form of catechol methylsideretin (*17*), confirmed by reduction (Supplementary Fig. S2, Note S2), and compounds *1* and *2* are non-canonical HCAs related to ferulic acid (see Supplementary Notes S2, S3 for further details).

All compounds were classified into coumarins (23) and HCAs (3) (Supplementary Fig. S4A). Most were detected across all genotypes, with only a few absent in Fe-inefficient lines (IsoClark and A97) (Supplementary Fig. S4B). Six compounds (23%) were exclusive to nutrient solutions (the two non-canonical HCAs, the oxidized forms of sideretin and catechol methylsideretin, and two trioxygenated coumarins, isomers of fraxetin and methylfraxetin), nine (35%) were found in both root extracts and nutrient solutions, including coumarin aglycones such as fraxetin, sideretin, and catechol methylsideretin, and eleven (42%) were only in root extracts (nine coumarin glycosides, ferulic acid, and one methylfraxetin isomer) (Supplementary Fig. S4C).

### Methylsideretin is the predominant phenolic compound released by soybean roots under iron deficiency

The 26 identified phenolics were quantified by HPLC–ESI–MS(TOF) using authenticated standards. None were detected at quantifiable levels in nutrient solutions from +Fe plants or in blanks; thus, quantification focused on nutrient solutions from −Fe plants sampled at Days 5 and 13, and on root extracts from both +Fe and −Fe plants at the same time points (Fig. 4).

**Fig. 4.**
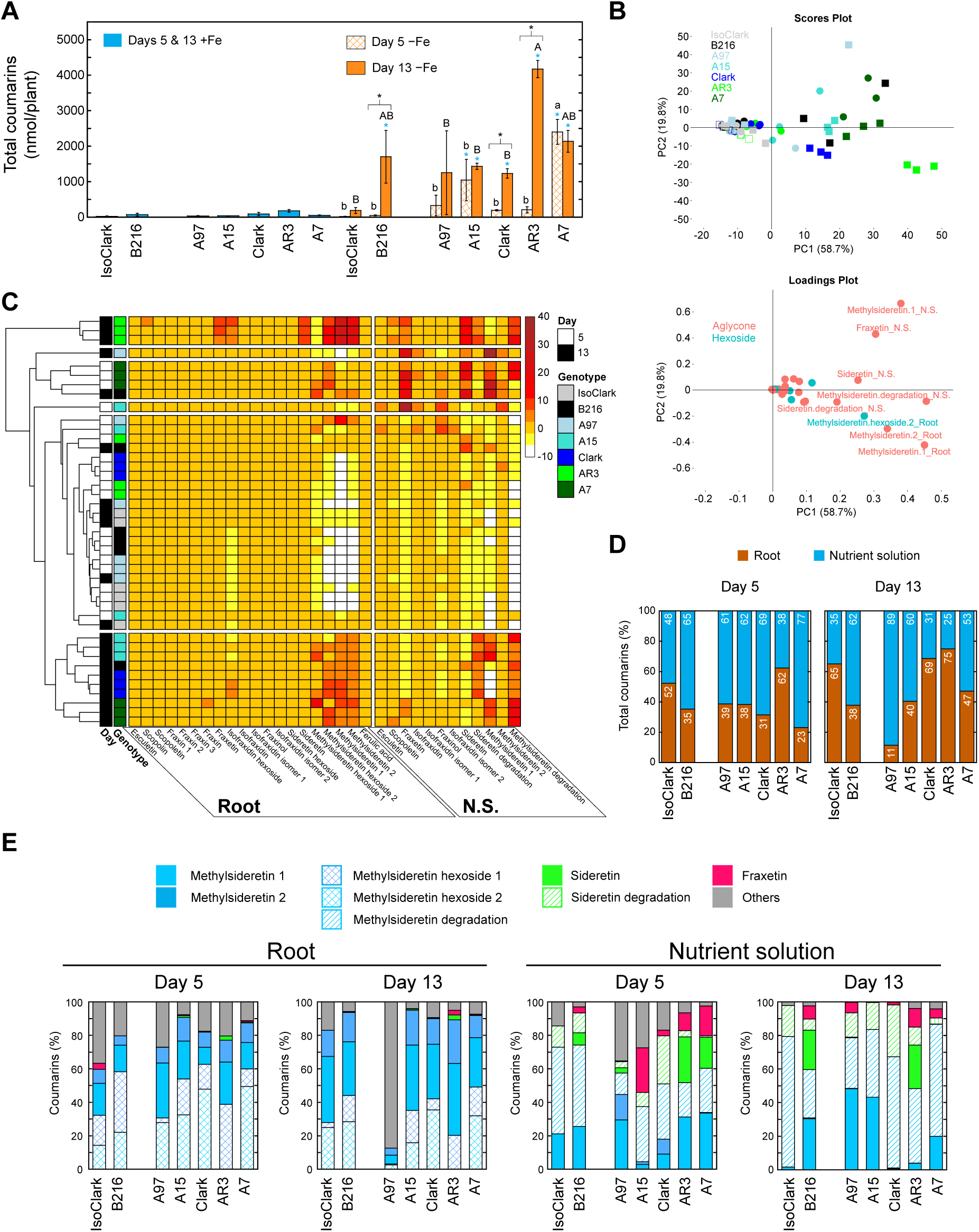
Quantitative profiles of the phenolic compounds root-accumulated and -released by seven soybean genotypes under iron deficiency. Plants were grown as described in Fig. 3. (A) Total phenolic content per plant (in nmol/plant). Columns represent means ± SE (n = 3 for −Fe at each sampling day; n = 4 for +Fe, combining Days 5 and 13). Significant differences between Fe treatments and sampling days within each genotype (*p* < 0.05) are indicated by blue and black asterisks, respectively. Under −Fe conditions, different lowercase (Day 5) and uppercase (Day 13) letters denote significant differences among genotypes (*p* < 0.05). (B) Principal component analysis of individual phenolic compound content. Scores plot: +Fe (empty symbols), −Fe (filled); Day 5 (circles), Day 13 (squares). Loadings plot: each compound is a filled circle. **C)** Hierarchical clustering heatmap of individual phenolic compound contents in root extracts and nutrient solutions (N. S.). (D) Distribution of phenolic compounds between roots and nutrient solutions under Fe deficiency. Data are means ± SE (n = 3). (E) Relative concentrations of phenolics in root extracts and nutrient solutions under Fe deficiency. Data are means ± SE (n = 3).

Total coumarin content per plant was calculated as the sum of all annotated coumarins present in roots and in the corresponding nutrient solutions. This calculation also included the two non-canonical HCAs, assumed to derive from degradation of sideretin and methylsideretin (see Supplementary Note S3 and Fig. S3A for details). Coumarin content varied among genotypes and was generally higher in −Fe plants than in +Fe ones (Fig. 4A). In +Fe roots, coumarin levels were low and similar across genotypes (23–178 nmol·plantL¹). On Day 5, only −Fe roots of two Fe-efficient genotypes, A7 and A15, showed significant increases compared with +Fe roots, with A7 (2399 nmol·plantL¹) accumulating more coumarins than A15 (1041 nmol·plantL¹). By Day 13, six of the seven genotypes produced significantly higher coumarin levels under −Fe conditions, the exception being the Fe-inefficient IsoClark. Among −Fe plants, AR3 reached the highest total coumarin content (4165 nmol·plantL¹), similar to A7 (2131 nmol·plantL¹), while the remaining genotypes showed lower values (1225–1693 nmol·plantL¹): the Fe-efficient A97, A15, and Clark, as well as the Fe-inefficient B216. Coumarin partitioning between roots and nutrient solution in −Fe plants showed genotype- and time-dependent patterns (Fig. 4D). At Day 5, five genotypes (A97, A15, Clark, A7, B216) allocated ≥60% of total coumarins to the nutrient solution, whereas IsoClark and AR3 allocated 45% and 38%, respectively. By Day 13, Clark secreted only ∼30% of total coumarins, A7 ∼53%, AR3 ∼25%, B216 ∼25%, and A97 increased secretion to 89%.

To examine variation in individual phenolic compounds, PCA was performed using the levels of all measured metabolites in roots and nutrient solutions (Fig. 4B). The first two principal components explained 79% of the variance (PC1, 59%; PC2, 20%). PC1 separated low-phenolic samples—including all +Fe and most Day 5 −Fe—from high-phenolic Day 13 −Fe samples, with A7 and A15 Day 5 −Fe samples clustering with the latter. PC2 captured variation among Day 13 −Fe samples, with AR3 forming a distinct subgroup. Loadings on PC1 were dominated by catechol methylsideretin, fraxetin, sideretin, and their degradation products in nutrient solutions, together with methylsideretin aglycones and the main hexoside in roots. Loadings on PC2 reflected differences in the distribution of these catechol coumarins between roots and nutrient solution, highlighting the distinctive metabolite profile of Day 13 −Fe AR3.

Hierarchical clustering of −Fe samples (Fig. 4C) indicated that the most abundant compounds—two methylsideretin isomers, sideretin, and fraxetin, including aglycone, glycoside, and degradation forms—largely determined sample grouping. Day 13 AR3 samples formed a distinct cluster with the highest levels in roots and nutrient solutions. Day 5 A7 and A15 samples, together with one Day 13 B216 and A97 sample, clustered separately with lower root fraxetin and fraxin. Day 13 A7, Clark, and A15, plus one Day 13 B216 sample, formed another cluster with similar methylsideretin and sideretin levels but low fraxetin. Remaining samples, mostly Day 5 −Fe, clustered together with the lowest levels of these four compounds.

We next examined the contribution of individual coumarin forms to total coumarin content in roots and nutrient solutions of −Fe plants (Fig. 4E). In roots, methylsideretin forms dominated across genotypes and sampling days, accounting for 60–95% of total root coumarins (except in A97 at Day 13). At Day 5, methylsideretin hexoside 2 was most abundant in A15, Clark, AR3, and A7, whereas at Day 13, catechol methylsideretin predominated in IsoClark, B216, A97, A15, and AR3. In nutrient solutions, catechol coumarins—including catechol methylsideretin, sideretin, their degradation products, and fraxetin—comprised 65–100% of total secreted coumarins, with either catechol methylsideretin or its degradation product predominating depending on genotype and sampling day. Together, these two compounds accounted for the majority of secreted coumarins, highlighting their dominant role in −Fe phenolic profiles.

### Coumarin secretion coincides with other physiological iron deficiency responses in iron-efficient soybean

To relate coumarin secretion to other Fe-deficiency responses, we analyzed the temporal dynamics of major phenolics (catechol methylsideretin, sideretin, fraxetin, and their degradation products) in the nutrient solutions of all genotypes. Here, we describe in detail the contrasting patterns of IsoClark and A7 shown in Figure 5 (with other genotypes presented in Supplementary Fig. S5).

**Fig. 5.**
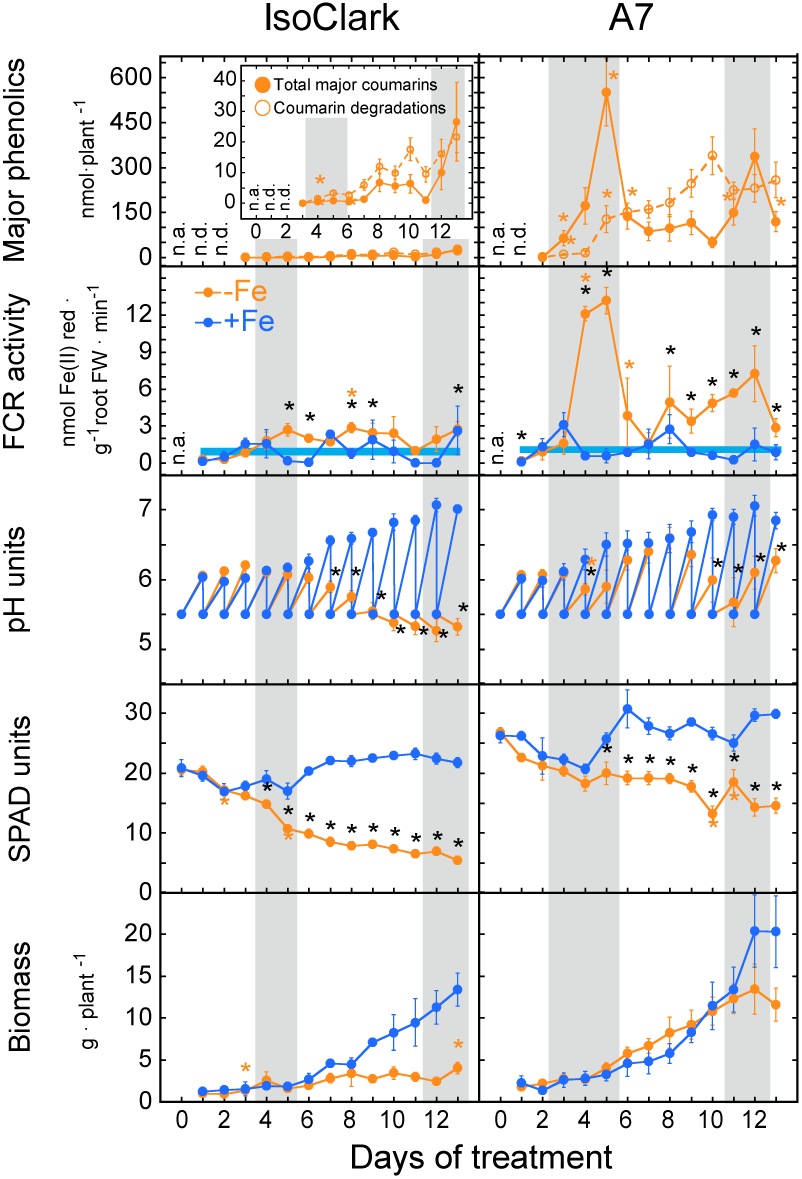
Temporal dynamics of root-released phenolics and key physiological responses to Fe deficiency in soybean genotypes with contrasting iron efficiency. Plants of IsoClark (Fe-inefficient) and A7 (Fe-efficient) were grown as described in Fig. 3. Time-course data are shown for phenolic compounds in the nutrient solution (NS), root ferric chelate reductase (FCR) activity, NS pH, SPAD values of young leaves, and plant fresh biomass in both genotypes. For phenolics in NS (only detected under −Fe), values represent the mean ± SE (n = 2–17, depending on genotype and day) of total content (nmol·plant^-1^) for two groups: major coumarins (catechol methylsideretin, sideretin, fraxetin) and putative degradation products of catechol methylsideretin and sideretin. IsoClark data are shown in a zoomed inset for clarity. ‘n.a.’ and ‘n.d.’ denote not available and not detected, respectively. For FCR, pH, SPAD, and biomass, data are shown for both −Fe and +Fe; sample sizes vary by parameter and treatment: FCR (n = 2 for +Fe; n = 3–4 for −Fe/day), NS pH (n = 4–11), and SPAD/biomass (n = 2–11). Significant differences between Fe treatments within each genotype and day (p < 0.05, t-test) are marked with a black asterisk above −Fe values. Orange asterisks indicate significant changes relative to the previous day in −Fe plants (*p* < 0.05, t-test). Grey shaded bands mark the time windows of phenolic secretion onset and peak, serving as temporal references for other physiological responses.

In both genotypes, major coumarins and their degradation products showed broadly similar temporal patterns in nutrient solution, yet genotypes differed markedly in level. IsoClark released only low amounts (<40 nmol·plant□¹), showing a modest rise at Day 4 followed by a gradual increase, including a late but non-significant elevation at Day 13. In contrast, A7 secreted much higher levels (>50 nmol·plant□¹ at most time points, exceeding 300 nmol·plant□¹ at Days 5, 10, and 12) and exhibited two pronounced secretion peaks around Days 5 and 11–12.

Other physiological responses also differed sharply between these two genotypes. IsoClark exhibited low, weakly inducible root FCR activity under −Fe conditions, with no clear temporal association with coumarin accumulation in nutrient solutions (Fig. 5). Medium acidification occurred mainly at Days 10–13, coinciding with pronounced chlorosis and biomass loss (Supplementary Fig. S6). In contrast, A7 displayed strong, recurrent FCR induction at nine time points, with peaks at Days 4–5 and 12 that coincided with major coumarin accumulation in nutrient solutions and marked medium acidification (Fig. 5). These synchronized responses were accompanied by slight regreening of −Fe young leaves at Day 11 (Fig. 5) and the first detectable decreases in biomass (Supplementary Fig. S6). Mineral analysis at Day 13 confirmed Fe depletion in all tissues and revealed a consistent Cu increase across organs (Supplementary Fig. S7).

### Iron deficiency strongly induces coumarin biosynthetic genes in Fe-efficient soybean roots

Transcript levels of soybean genes homologues of Arabidopsis genes involved in coumarin biosynthesis, transport, and classical Fe-uptake strategy genes (*FER-like*, *FRO2*, *IRT* and *AHA4*) were measured by RT-qPCR in roots of −Fe and +Fe plants at Days 3–5. For simplicity, Figure 6 shows −Fe Day 4 and +Fe Days 3–5, as most +Fe roots showed little temporal variation (Supplementary Figs. S8–S10). Among classical Fe-uptake genes, only *GmIRT-like* was significantly induced in A7 (∼22-fold), whereas *GmFER-like*, *GmFRO2*, and *GmAHA4* showed modest, non-significant changes (Fig. 6 top panel). In IsoClark, none of these genes were significantly induced.

**Fig. 6.**
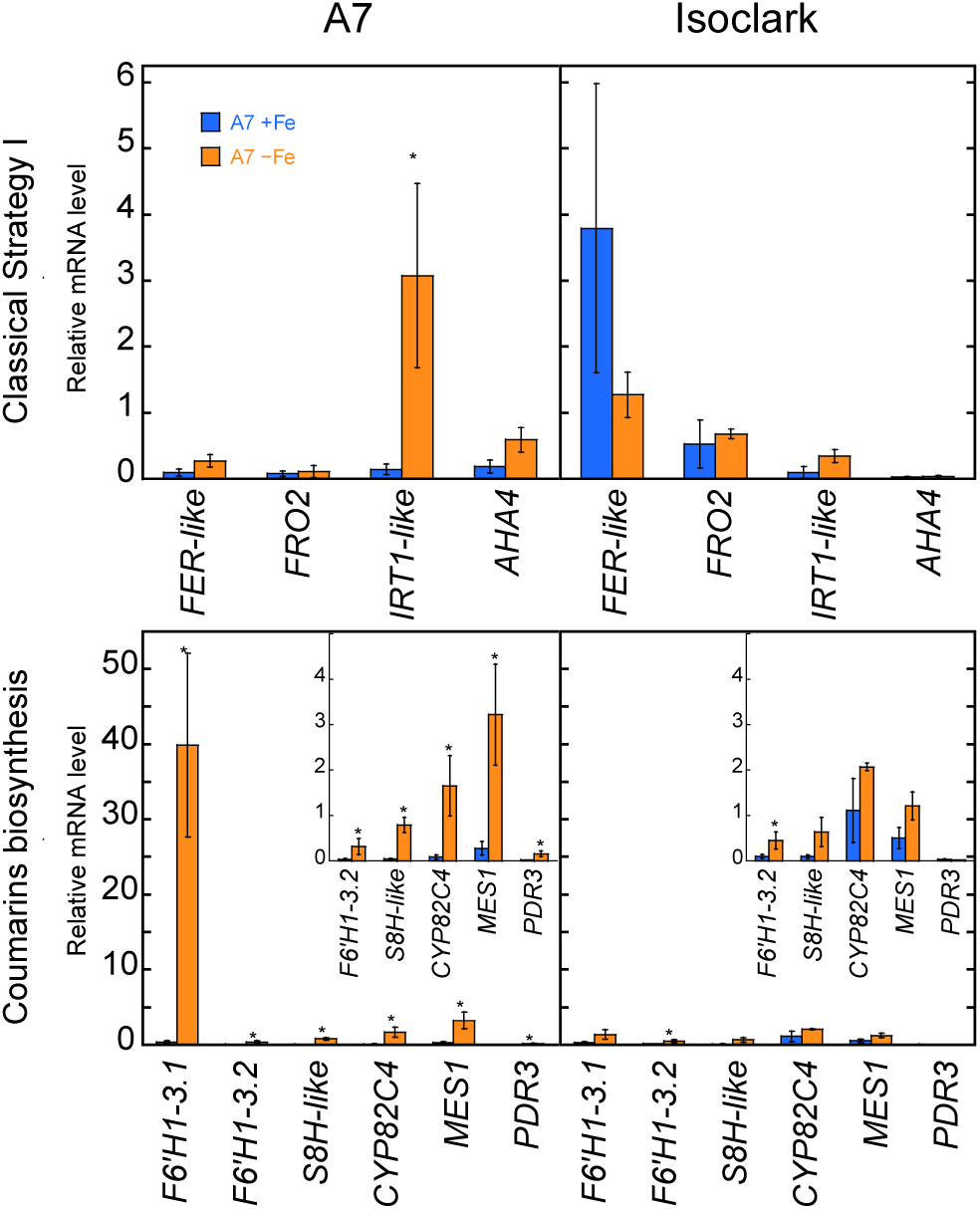
RT-qPCR of key iron deficiency-related genes in soybean genotypes with contrasting iron efficiency. Plants of IsoClark (Fe-inefficient) and A7 (Fe-efficient) were grown as described in Fig. 3. Roots were sampled on Days 3, 4, and 5 of treatment from Fe-sufficient (+Fe) and Fe-deficient (−Fe) plants. Transcript levels are shown for classical Fe-deficiency response genes of the reduction-based strategy (*FER*, *FRO2*, *IRT1*, *AHA4*), genes of the phenylpropanoid/coumarin-related pathway (*F6’H1-3.1*, *F6’H1-3.2*, *S8H-like*, *CYP82C4)*, a putative methyl-transferase (*MES1*), and a putative coumarins transporter (*PDR3*, the soybean homolog of *AtPDR9*). For −Fe plants, transcript levels correspond to Day 4; for +Fe plants, to the mean of Days 3–5. Relative transcript levels were calculated as Normalized Relative Quantity (NRQ) according to Hellemans et al. (2007). Data are means ± SE (n = 3-6). Significant differences between Fe treatments for each and genotype (*p* < 0.05) are indicated by an asterisk above the −Fe column.

Phylogenetic analysis of soybean ABCG transporters identified two candidates closely related to Arabidopsis AtPDR9/AtABCG37 and tobacco NtPDR3/NtABCG3 (Supplementary Fig. S11). Of these, only *GmPDR3* (*GLYMA_17G039300*) was induced by Fe deficiency in A7 (∼7-fold) (Fig. 6 bottom panel), consistent with a role as coumarin exporter.

Genes of the coumarin biosynthetic pathway, including *GmF6’H1-3.1* (*GLYMA_07G124400*), *GmF6’H1-3.2* (*GLYMA_03G096500*), *GmS8H-like* (*GLYMA_08G169100*), and *GmCYP82C4* (*GLYMA_04G035600*), were all strongly induced in A7. *GmF6’H1-3.1* showed the highest increase (∼120-fold), *GmS8H-like* and *GmCYP82C4* up to 20-fold, and *GmF6’H1-3.* ∼12-fold (Fig. 6 bottom panel). Methyltransferase analysis revealed strong induction of *GmMES1* (∼12-fold), a putative enzyme for methylsideretin production, whereas homologues of AtOMT1 showed no significant changes. In IsoClark, only *GmF6’H1-3.2* was induced, and to a lower extent (∼5-fold).

### Catechol methylsideretin in soybean root exudates correlates with their ability to solubilize ferric oxide

To assess the contribution of individual coumarins to the Fe mobilization rate found for plant nutrient solutions of −Fe plants, coumarin concentrations in nutrient solutions for each genotype were quantified by HPLC-ESI-MS(TOF) (Supplementary Fig. S12). Correlations between coumarin levels and total Fe mobilized at pH 5.5 and 7.5 were evaluated, considering total coumarins, total coumarins plus degradation products, the catechol subgroup, and individual catechol coumarins (catechol methylsideretin, fraxetin, and sideretin) (Fig. 7A).

**Fig. 7.**
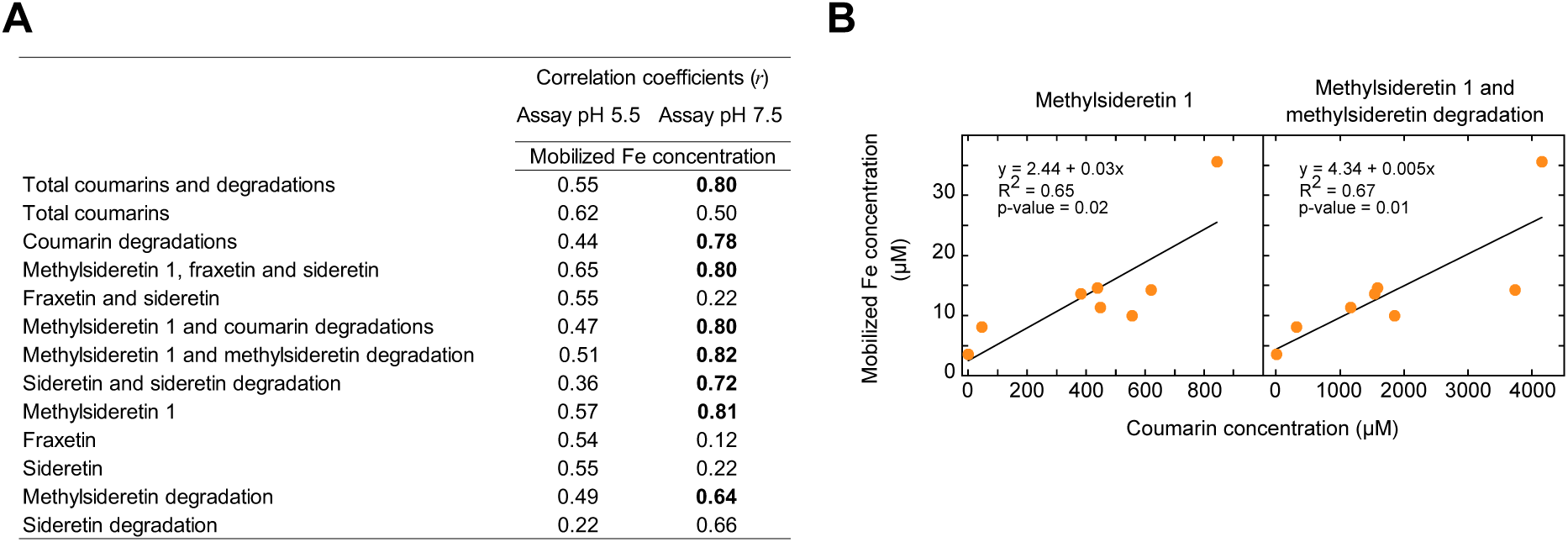
Correlation between phenolic compound concentrations and iron mobilized from ferric oxide. Pools of nutrient solutions from each genotype used in the Fe mobilization assay (Figure 2) were analyzed for phenolics by HPLC-ESI-MS(TOF). (A) Correlation coefficients between the concentrations (µM) of the phenolic compounds or compound groups in the assay solution and the concentration of Fe mobilized (µM) after 6 h incubation with ferric oxide in the presence of 600 µM BPDS and 1 mM NADH at pH 5.5 and 7.5. (B) Scatter plots showing the concentration of catechol methylsideretin (µM), or catechol methylsideretin plus its degradation product (µM), versus the concentration of Fe mobilized (µM) at pH 7.5. Linear regression lines are shown, along with equations, coefficients of determination (R²), and significance levels (*p*-values).

No significant correlations were observed at pH 5.5. At pH 7.5, Fe mobilization correlated strongly with catechol methylsideretin and its degradation product (r = 0.82) and with catechol methylsideretin alone (r = 0.81). The combined total coumarins plus degradation products also showed a significant correlation (r = 0.80). Linear regression confirmed a moderate-to-strong association between catechol methylsideretin and Fe mobilization, slightly enhanced when including its degradation products (Fig. 7B).

## DISCUSSION

We provide the first detailed chemical characterization of Fe deficiency–induced compounds released by soybean roots. Among nearly two dozen coumarins, a previously undescribed catechol-type methylsideretin was the dominant component in roots and nutrient solutions (Fig. 4E), while the remaining identified compounds have been previously reported in *A. thaliana* and tobacco. Using seven genotypes with contrasting Fe efficiency, we linked variation in coumarin production and secretion to differences in Fe mobilization from ferric oxide at circumneutral pH (Fig. 2). Production and release of these compounds increased under Fe deficiency, particularly in IDC-tolerant genotypes, coinciding with other root traits involved in efficient Fe acquisition (Fig. 5). Consistent with this, the coumarin biosynthetic pathway was upregulated in Fe-efficient, but not Fe-inefficient, roots (Fig. 6). Coumarin profiles were qualitatively similar across genotypes (Supplementary Fig. S4), with the most quantitative variation attributable to catechol methylsideretin (Fig. 4A,C), which also showed the strongest correlation with Fe mobilization (Fig. 7). Soybean remains the only legume with chemically defined phenolic exudates under Fe deficiency; other legumes such as peanut, pea, or red clover release phenolics under Fe deficiency (Römheld and Marschner 1983; Jelali et al., 2010; Jin et al., 2007) but they remain uncharacterized, whereas *Medicago truncatula* and *M. scutellata* secrete flavins instead (Rodríguez-Celma et al., 2011; Gheshlaghi et al., 2021).

Coumarins accounted for 23 of the 26 phenolic compounds consistently detected in roots and nutrient solutions of Fe-deficient soybean genotypes (Tables 1, Supplementary Table S4 and Fig. S4). Most were highly oxygenated, a feature known to favor exudation via the Fe deficiency–induced PDR9 transporter (Ziegler et al., 2018). Catechol methylsideretin, identified here for the first time as Fe-deficiency–elicited, is structurally similar to sideretin with a critical catechol moiety for Fe mobilization (Schmid et al., 2014; Sisó-Terraza et al., 2016a; Rajniak et al., 2018). Both oxidized and reduced forms were detected, with the oxidized form restricted to nutrient solutions, reflecting facile oxidation under aerated hydroponic conditions (Supplementary Fig. S4; Rajniak et al., 2018; Lefèvre et al., 2018). The redox potential (E°) of oxygenated coumarins is largely determined by the substituent at the para position relative to the catechol moiety (Liu et al., 2014). Sideretin carries a hydroxyl group (-OH), which increases its redox potential and facilitates oxidation, explaining its predominance in the oxidized form in nutrient solutions (Rajniak et al., 2018; Lefèvre et al., 2018; Ziegler et al., 2018). Fraxetin lacks a para substituent (-H), resulting in lower redox potential and slower oxidation, while catechol methylsideretin carries a methoxy group (-OCH_3_), likely conferring an intermediate redox potential and oxidation behavior between sideretin and fraxetin. These differences underlie their distinct ecological roles: sideretin primarily acts as a strong reductant favoring Fe²L uptake at acidic pH, whereas fraxetin mobilizes Fe³L via complexation, which is more relevant at neutral to alkaline pH (Paffrath et al., 2024). Catechol methylsideretin may combine both reductive and chelating functions, potentially complementing the activities of sideretin and fraxetin in Fe acquisition under variable soil conditions, though its redox behavior and ecological function require experimental validation.

The remaining coumarins identified in the present work, including sideretin, fraxetin and a different methylsideretin isomer, have been reported in *A. thaliana* and tobacco (Supplementary Table S6). Fourteen coumarins were found as aglycones, while glycosides were exclusively root-localized, consistent with prior observations in *A. thaliana* and tobacco (Fourcroy et al., 2014; Sisó-Terraza et al., 2016a; Lefèvre et al., 2018), supporting a role for glycosylation/de-glycosylation in coumarin storage and transport (Robe et al., 2021).

Besides coumarins, three other phenolic compounds—HCAs—were detected. One of these, ferulic acid, serves as a precursor in coumarin biosynthesis: it is converted to feruloyl-CoA, which enters the first step of scopoletin formation (Fig. 1), from which the more complex coumarins detected in soybean roots are likely derived, as reported for fraxetin and sideretin in *A. thaliana* (Tsai et al., 2018; Rajniak et al., 2018). Ferulic acid was found only as an aglycone in roots (Supplementary Fig. S7), whereas in *A. thaliana* roots its glycoside form accumulates under Fe deficiency (Schmid et al., 2014; Sisó-Terraza et al., 2016a). The other two HCAs were non-canonical, detected exclusively in nutrient solutions, and tentatively annotated as degradation products of catechol methylsideretin and sideretin, based on MS^n^ fragmentation patterns (Supplementary Notes S2, S3; Fig. S3). These annotations are consistent with reports that highly substituted coumarins are more prone to degradation (Kang et al., 2023), but remain tentative in the absence of commercially available authenticated standards.

This study provides the most comprehensive intraspecific analysis of Fe deficiency–induced coumarin profiles in plants to date. Previous work in *A. thaliana* either quantified a single coumarin (fraxetin) across 22 accessions with variable Fe responses (Tsai et al., 2018) or profiled coumarins only in two small local populations with contrasting tolerance to soil carbonate (Terés et al., 2018), but none addressed genotype-dependent variation at this depth. The seven soybean genotypes examined under Fe deficiency exhibited qualitatively similar coumarin profiles, with all genotypes sharing the same set of compounds (Supplementary Fig. S4). In an interspecific comparison, *A. thaliana*, tobacco, and soybean share a largely similar core set of coumarins, including sideretin, fraxetin, scopoletin, isofraxidin, and fraxinol. Soybean and *A. thaliana* additionally produce esculetin and isofraxetin, soybean and tobacco synthesize methylsideretin, and uniquely, only soybean accumulates catechol methylsideretin. In contrast, other plant species exhibit simpler coumarin profiles under Fe deficiency: dihydrocoumarin and xanthotoxin in tomato (Astolfi et al., 2020), 4-hydroxycoumarin in *Vitis champini* (Marastoni et al., 2020), and limited sets in *Brassica rapa* (sideretin), *Eutrema salsugineum* (esculetin, isoscopoletin, methylated 5,6,7-trihydroxycoumarins), and *Leucanthemum vulgare* (sideretin) (Rajniak et al., 2018). These patterns highlight the diversity of coumarin secretion across species. However, many studies assessed coumarins only as a minor component of broader analyses, so analytical methods, growth conditions, and sampling time likely affect the apparent diversity. Future targeted investigations could reveal a broader coumarin spectrum in these species.

Significant genotypic differences in total coumarin production were observed among soybean genotypes with contrasting Fe efficiencies, primarily in terms of magnitude and timing. Genotypes ranged from low producers (Fe-inefficient IsoClark and B216; Fe-efficient A97, which showed high replicate-to-replicate variability leading to a relatively low mean) to intermediate producers (Clark, A15) and high producers (A7, AR3) (Figs. 4A,B, 5 and Supplementary Fig. S5). Timing was critical: some genotypes initiated coumarin production early under Fe deficiency (strongly in A7, moderately in A15), others showed delayed but sometimes greater accumulation (AR3), and some exhibited moderate late-stage increases (B216, A97, Clark). A7 and A15 not only initiated synthesis earlier but also displayed repeated secretion events, indicating sustained metabolic adaptation. Common breeding parentage may partly explain these patterns: A7 and A15 share the AP9 lineage, AR3 derives from a cross with the variable producer A97, while B216 is more distantly related (Jolley et al., 1986; Bernard et al., 1991; Peiffer et al., 2012). In contrast, Clark and its near-isogenic line IsoClark, which carries a 12-bp deletion in the bHLH38 transcription factor (Peiffer et al., 2012), differ sharply in coumarin production and Fe efficiency. This deletion impairs FIT activity, required in *A. thaliana* for transcriptional regulation of coumarin biosynthesis enzymes F6’H1, S8H, and CYP82C4 under Fe deficiency (Schmid et al., 2014; Rajniak et al., 2018). Since FIT functions as a heterodimer with bHLH38/39, loss of bHLH38 reduces FIT activity and coumarin production, explaining IsoClark’s weak but temporally similar secretion pattern. Supporting this, gene expression data (Fig. 6, Supplementary Figs. S8, S9) show strong upregulation of coumarin biosynthesis genes in A7 under Fe deficiency, whereas IsoClark shows no induction, consistent with previous reports (Peiffer et al., 2012; Waters et al., 2018). Moreover, since ectopic expression of AtbHLH38/39 in other dicotyledonous plants such as tobacco induces flavin synthesis and secretion even under Fe sufficiency (Vorwieger et al., 2007), observations in IsoClark and A7 support a role for bHLH38 in controlling coumarin induction under Fe deficiency. These findings suggest that targeting bHLH38-FIT regulation could guide breeding strategies to enhance soybean Fe efficiency.

Genotypic differences exist in the partitioning of coumarins between roots and the growth medium in soybean. This effect was much smaller than the variation observed in total coumarin production under Fe deficiency (Fig. 4A,D). Most genotypes accumulated between 53% and 89% of total coumarins in the nutrient solution, with the remainder retained in roots. In contrast, the Fe-inefficient IsoClark and the Fe-efficient AR3 showed markedly lower secretion ratios (35–48% and 25–38%, respectively), suggesting some impairment in export. This pattern resembles that observed in *Atpdr9* mutants and NtPDR3-silenced tobacco lines, where loss of PDR transporter function leads to intracellular coumarin accumulation and reduced secretion (Fourcroy et al., 2014; Ziegler et al., 2017; Lefèvre et al., 2018). Similar effects have been reported in Arabidopsis mutants defective in BETA-GLUCOSIDASE 42 (*bglu42*), which are unable of scopolin deglycosylation, limiting scopoletin release from root hairs under Fe deficiency (Zamioudis et al., 2014; Robe et al., 2021). Therefore, the altered coumarin distribution in IsoClark and AR3 may result from defective transporter activity, impaired deglycosylation, or both. In IsoClark, the mutation in *bHLH38* could also affect genes involved in export, while in AR3, the low secretion despite high internal accumulation suggests a persistent transport limitation associated with its delayed yet abrupt biosynthetic induction (Fig. 4D, Supplementary Fig. S8).

Methylsideretin forms predominated in soybean under Fe deficiency, accounting for >60% of total coumarins in roots and >40% in nutrient solutions across all genotypes and sampling times (Fig. 4E). Within roots, all methylsideretin forms (aglycone isomers and glycosides) accounted for 60–95% of total coumarins, with glycosides dominating. Nutrient solutions had greater diversity, enriched in catechol coumarins; catechol methylsideretin aglycone and its degradation product typically exceeded 50% of total coumarins, with the remainder including sideretin, its degradation product, and fraxetin. This resembles Fe-deficient tobacco roots, where methylsideretin glycosides predominated in roots, despite only one aglycone isomer being detected in nutrient solutions (Lefèvre et al., 2018). By contrast, *A. thaliana* accumulates scopolin as the major root coumarin, while exudates are dominated by sideretin and fraxetin (Rajniak et al., 2018; Paffrath et al., 2024). Genotype has little effect on relative coumarin composition. Correlation analyses showed catechol methylsideretin had the strongest positive association with Fe mobilization, strengthened when coumarin degradation products were included (Fig. 7A), and linear regression confirmed a moderate-to-strong relationship (Fig. 7B), supporting its key role in Fe solubilization, consistent with previous observations for the catechol coumarin fraxetin (Sisó-Terraza et al., 2016a). Beyond Fe mobilization, coumarins influence on root microbiomes: in *A. thaliana*, fraxetin and scopoletin modulate rhizosphere bacterial communities, selectively inhibiting certain bacterial taxa such as Pseudomonas species via redox-mediated mechanisms (Voges et al., 2019). Moreover, plant-microbe interactions mediated by redox-active metabolites vary with environmental conditions, including oxygen, pH, and exudate carbon sources, affecting microbial growth under Fe limitation (McRose et al., 2023). As catechol methylsideretin is a dominant exudate in soybean, genotypic variation could provide a model to study its impact on the root microbiome.

## SUPPLEMENTARY DATA

**Note S1.** Optimization of soybean growth conditions

**Note S2.** Detailed information on the annotation of compounds accumulated in nutrient solutions and root extracts of iron-deficient soybean plants

**Note S3.** Evidence supporting the annotation of the non-canonical HCAs (compounds 1 and 2) **Fig. S1.** Effects of genotype additives on Fe mobilization from ferric oxide by root exudates of seven soybean genotypes grown under iron deficiency

**Fig. S2.** Annotation of compounds 7 and 16

**Fig. S3.** Proposed origin of compounds 1 and 2 detected in the nutrient solution

**Fig. S4.** Occurrence of the annotated phenolic compounds in nutrient solutions and root extracts across soybean genotypes under iron deficiency

**Fig. S5.** Temporal dynamics of root-released phenolics and key physiological responses to Fe deficiency in soybean genotypes B216, A97, A15, Clark, and AR3

**Fig. S6.** Effects of iron deficiency on soybean biomass

**Fig. S7.** Effects of iron deficiency on micronutrient levels in different organs of soybean genotype A7

**Fig. S8.** Time-course analysis of Fe deficiency-related gene expression in roots of genotype A7 **Fig. S9.** Time-course analysis of Fe deficiency-related gene expression in roots of genotype IsoClark

**Fig. S10.** Time-course expression of a putative coumarin transporter gene in roots of A7 and IsoClark

**Fig. S11.** Phylogenetic tree of PDR family proteins from soybean, tobacco, and Arabidopsis thaliana

**Fig. S12.** Concentrations of phenolic compounds in nutrient solution pools used for iron mobilization assays

**Table S1.** HPLC elution program used for the separation of phenolic compounds and flavins

**Table S2.** HCT Ultra ion trap mass spectrometer settings

**Table S3.** Primers used for qPCR analysis

**Table S4.** MSⁿ fragmentation data for phenolic compounds released and accumulated by soybean roots under iron deficiency.

**Table S5.** Retention times and TOF/Ion Trap mass spectrometric data for authenticated phenolic standards used for compound annotation.

**Table S6.** Coumarin-type phenolics released and/or accumulated by soybean and other plant species in response to iron deficiency

## ACKNOWLEDGEMENTS

We thank Gema Marco, Leticia Berlanga, and Margarita Palancar (Estación Experimental de Aula Dei, EEAD-CSIC) for their invaluable technical assistance with plant handling, phenolic compound quantification, and mineral analysis, respectively.

## AUTHOR CONTRIBUTIONS

AA-F and EG-C conceived and designed the experiments. EG-C conducted the plant experiments and collected the corresponding data. FJJ-P and EG-C identified and quantified phenolic compounds. FJJ-P performed the iron mobilization, processed the data, prepared figures, and drafted the manuscript. FJJ-P and RB performed gene expression analyses. AA-F supervised the plant experiments, phenolic compound analyses, and iron mobilization assays. JR-C supervised the gene expression work. JA, JR-C, and AA-F contributed to writing, reviewing, and editing the manuscript and secured the funding.

## DATA AVAILABILITY

The metabolomics MS data supporting the findings of this study have been deposited in the MassIVE repository under dataset identifier MSV000101246. Additional data are available from the corresponding author, Ana Álvarez-Fernández, upon reasonable request

## ABBREVIATIONS

ABCG37: ATP-BINDING CASSETTE G37
BGLU42: BETA-GLUCOSIDASE 42
CID: collision-induced dissociation
COSY: COUMARIN SYNTHASE
ESI: electrospray ionization
F6’H1: FERULOYL-CoA 6’-HYDROXYLASE 1
FCR: ferric chelate reductase
FIT: FER-LIKE IRON DEFICIENCY-INDUCED TRANSCRIPTION FACTOR
FRO2: FERRIC REDUCTION OXIDASE 2
HCAs: hydroxycinnamic acids
IDC: iron deficiency chlorosis
IRT1: IRON-REGULATED TRANSPORTER 1
ISs: internal standards
MA: mugineic acid
MS: mass spectrometry
NRQ: normalized relative quantity
PDR9: PLEIOTROPIC DRUG RESISTANCE 9
PM: plasma membrane
RTs: retention times
S8H: SCOPOLETIN 8-HYDROXYLASE
TOF: time-of-flight
UV/VIS: ultraviolet–visible spectrophotometry

## REFERENCES

1. Astolfi S, Pii Y, Mimmo T, Lucini L, Miras-Moreno MB, Coppa E, Violino S, Celletti S, Cesco S. 2020. Single and combined Fe and S deficiency differentially modulate root exudate composition in tomato: a double strategy for Fe acquisition? International Journal of Molecular Sciences 21: 4038.

2. Bernard RL, Nelson RL, Cremeens CR. 1991. USDA soybean genetic collection: isoline collection. Soybean Genetics Newsletter 18: 27–57.

3. Bienfait HF, Bino RJ, Vanderbliek AM, Duivenvoorden JF, Fontaine JM. 1983. Characterization of ferric reducing activity in roots of Fe-deficient *Phaseolus vulgaris*. Physiologia Plantarum 59: 196–202.

4. Brown JC, Holmes RS, Tiffin L. 1961. Iron chlorosis in soybeans as related to the genotype of rootstalk: Chlorosis susceptibility and reductive capacity at the root. Soil Science 91: 127–132.

5. Brown JC, Ambler JE. 1973. “Reductants” Released by Roots of Fe-Deficient Soybeans. Agronomy Journal 65: 311–314.

6. Borges F, Roleira F, Milhazes N, Santana L, Uriarte E. 2005. Simple coumarins and analogues in medicinal chemistry: occurrence, synthesis and biological activity. Current Medicinal Chemistry 12: 887–916.

7. Fourcroy P, Sisó-Terraza P, Sudre D, Savirón M, Reyt G, Gaymard F, Abadía A, Abadía J, Álvarez-Fernández A, Briat JF. 2014. Involvement of the ABCG37 transporter in secretion of scopoletin and derivatives by Arabidopsis roots in response to iron deficiency. New Phytologist 201: 155–167.

8. Gheshlaghi Z, Luis-Villarroya A, Álvarez-Fernández A, Khorassani R, Abadía J. 2021. Iron deficient *Medicago scutellata* grown in nutrient solution at high pH accumulates and secretes large amounts of flavins. Plant Science 303: 110664.

9. González-Guerrero M, Navarro-Gómez C, Rosa-Núñez E, Echávarri-Erasun C, Imperial J, Escudero V. 2023. Forging a symbiosis: transition metal delivery in symbiotic nitrogen fixation. New Phytologist 239: 2113–2125.

10. Hellemans J, Mortier G, De Paepe A, Speleman F, Vandesompele J. 2007. qBase relative quantification framework and software for management and automated analysis of real-time quantitative PCR data. Genome Biology 8: R19.

11. Herridge DF, Peoples MB, Boddey RM. 2008. Global inputs of biological nitrogen fixation in agricultural systems. Plant and Soil 311: 1–18.

12. Jelali N, M’sehli W, Dell’Orto M, Abdelly C, Gharsalli M, Zocchi G. 2010. Changes of metabolic responses to direct and induced Fe deficiency of two *Pisum sativum* cultivars. Environmental and Experimental Botany 68: 238–246.

13. Jin CW, You GY, He YF, Tang CX, Wu P, Zheng SJ. 2007. Iron deficiency-induced secretion of phenolics facilitates the reutilization of root apoplastic iron in red clover. Plant Physiology 144: 278–285.

14. Jolley VD, Brown JC, Davis TD, Walser RH. 1986. Increased Fe-efficiency in soybean through plant breeding related to increased response to Fe-deficiency stress. I. Iron stress response. Journal Plant Nutrition 9: 373–386.

15. Kang K, Schenkeveld WDC, Weber G, Kraemer SM. 2023. Stability of coumarins and determination of the net iron oxidation state of iron-coumarin complexes: implications for examining plant iron acquisition mechanisms. ACS Earth Space Chemistry 7: 2339–2352.

16. Kobayashi T, Nishizawa NK. 2012. Iron Uptake, Translocation, and Regulation in Higher Plants. Annual Review of Plant Biology 63: 131–152.

17. Lefèvre F, Fourmeau J, Pottier M, Baijot A, Cornet T, Abadía J, Álvarez-Fernández A, Boutry M. 2018. The Nicotiana tabacum ABC transporter NtPDR3 secretes O-methylated coumarins in response to iron deficiency. Journal of Experimental Botany 69: 4419–4431.

18. Lindsay WL. 1995. Chemical reactions in soils that affect iron availability to plants. A quantative approach. In: Abadía J, ed. Iron Nutrition in Soils and Plants. Dordrecht, the Netherlands: Kluwer Academic, 7–14.

19. Liu T, Liu MM, Zheng XW, Du CY, Cui XY, Wang L, Han LL, Yu ZY. 2014. Substituent effects on the redox potentials of dihydroxybenzenes: theoretical and experimental study. Tetrahedron 70: 9033–9040.

20. McRose DL, Li J, Newman DK. 2023. The chemical ecology of coumarins and phenazines affects iron acquisition by pseudomonads. Proceedings of the National Academy of Sciences, USA 120: e2217951120.

21. Merry R, Dobbels AA, Sadok W, Naev S, Stupar RM, Lorenz AJ. 2022. Iron deficiency in soybean. Crop Science 62: 36–52.

22. Moran-Lauter AN, Peiffer GA, Yin T, Whitham SA, Cook D, Shoemaker RC, Graham MA. 2014. Identification of candidate genes involved in early iron deficiency chlorosis signaling in soybean (*Glycine max*) roots and leaves. BMC Genomics 15: 702.

23. Paffrath V, Tandron Moya YA, Weber G, von Wirén N, Giehl RFH. 2024. A major role of coumarin-dependent ferric iron reduction in strategy I-type iron acquisition in Arabidopsis. Plant Cell 36: 642–664.

24. Peiffer GA, King KE, Severin AJ, May GD, Cianzio SR, Lin SF, Lauter NC, Shoemaker RC. 2012. Identification of candidate genes underlying an iron efficiency quantitative trait locus in soybean. Plant Physiology 158: 1745–54.

25. Raj KK, Pandey RN, Singh B, Talukdar A. 2019. ^14^C labelling as a reliable technique to screen soybean genotypes (*Glycine max* (L.) Merr.) for iron deficiency tolerance. Journal of Radioanalytical and Nuclear Chemistry 322: 655–662.

26. Rajniak J, Giehl RFH, Chang E, Murgia I, von Wirén N, Sattely ES. 2018. Biosynthesis of redox-active metabolites in response to iron deficiency in plants. Nature Chemical Biology 14: 442–450.

27. Ren Z, Zhang L, Li H, et al. 2025. The BRUTUS iron sensor and E3 ligase facilitates soybean root nodulation by monoubiquitination of NSP1. Nature Plants 11: 595–611.

28. Robinson N, Procter C, Connolly E. 1999. A ferric-chelate reductase for iron uptake from soils. Nature 397: 694–697.

29. Robe K, Conejero G, Gao F, et al. 2021. Coumarin accumulation and trafficking in Arabidopsis thaliana: a complex and dynamic process. New Phytologist 229: 2062–2079.

30. Robe K, Stassen MJJ, Watanabe S, et al. 2025. Coumarin-facilitated iron transport: An IRT1-independent strategy for iron acquisition in *Arabidopsis thaliana*. Plant Communications 6: 101431.

31. Rodríguez-Celma J, Vázquez-Reina S, Orduna J, Abadía A, Abadía J, Álvarez-Fernández A, López-Millán AF. 2011. Characterization of flavins in roots of Fe-deficient strategy I plants, with a focus on *Medicago truncatula*. Plant Cell Physiology 52: 2173–89.

32. Rodríguez-Celma J, Lin WD, Fu GM, Abadía J, López-Millán AF, Schmidt W. 2013. Mutually exclusive alterations in secondary metabolism are critical for the uptake of insoluble iron compounds by Arabidopsis and *Medicago truncatula*. Plant Physiology 162: 1473–1485.

33. Römheld V, Marschner H. 1983. Mechanism of iron uptake by peanut plants I. Fe^III^ reduction, chelate slitting, and release of phenolics. Plant Physiology 71: 949–954.

34. Römheld V, Marschner H. 1986. Evidence for a specific uptake system for iron phytosiderophores in roots of grasses. Plant Physiology 80: 175–180.

35. Santi S, Schmidt W. 2009. Dissecting iron deficiency-induced proton extrusion in Arabidopsis roots. New Phytologist 183: 1072–1084.

36. Schmid NB, Giehl RF, Döll S, Mock HP, Strehmel N, Scheel D, Kong X, Hider RC, von Wirén N. 2014. Feruloyl-CoA 6’-Hydroxylase1-dependent coumarins mediate iron acquisition from alkaline substrates in Arabidopsis. Plant Physiology 164: 160–172.

37. Schmidt H, Günther C, Weber M, 2014. Metabolome analysis of *Arabidopsis thaliana* roots identifies a key metabolic pathway for iron acquisition. PloS One 9: e102444.

38. Sisó-Terraza P, Luis-Villarroya A, Fourcroy P, Briat JF, Abadía A, Gaymard F, Abadía J, Álvarez-Fernández A. 2016a. Accumulation and secretion of coumarinolignans and other coumarins by *Arabidopsis thaliana* roots in response to iron deficiency at high pH. Frontiners in Plant Science 7: 1711.

39. Sisó-Terraza P, Rios JJ, Abadía J, Abadía A, Álvarez-Fernández A. 2016b. Flavins secreted by roots of iron-deficient *Beta vulgaris* enable mining of ferric oxide via reductive mechanisms. New Phytologist 209: 733–45.

40. Terés J, Busoms S, Perez Martín L, Luís-Villarroya A, Flis P, Álvarez-Fernández A, Tolrà R, Salt DE, Poschenrieder C. 2019. Soil carbonate drives local adaptation in *Arabidopsis thaliana*. Plant, Cell and Environment 42: 2384–2398.

41. Tsai HH, Rodríguez-Celma J, Lan P, Wu YC, Vélez-Bermúdez IC, Schmidt W. 2018. Scopoletin 8-hydroxylase-mediated fraxetin production is crucial for iron mobilization. Plant Physiology 177: 194–207.

42. Usman M, Li Q, Luo D, Xing Y, Dong D. 2025. Valorization of soybean by-products for sustainable waste processing with health benefits. Journal of Science and Food Agriculture 105: 5150–5162.

43. Vanholme R, Sundin L, Seetso KC, et al. 2019. COSY catalyses trans-cis isomerization and lactonization in the biosynthesis of coumarins. Nature Plants 5: 1066–1075.

44. Vert G, Grotz N, Dédaldéchamp F, Gaymard F, Guerinot ML, Briat JF, Curie C. 2002. IRT1, an Arabidopsis transporter essential for iron uptake from the soil and for plant growth. Plant Cell 14: 1223–1233.

45. Voges MJEEE, Bai Y, Schulze-Lefert P, Sattely ES. 2019. Plant-derived coumarins shape the composition of an Arabidopsis synthetic root microbiome. Proceedings of the National Academy of Sciences, USA 116: 12558–12565.

46. Vorwieger A, Gryczka C, Czihal A, Douchkov D, Tiedemann J, Mock HP, Jakoby M, Weisshaar B, Saalbach I, Bäumlein H. 2007. Iron assimilation and transcription factor controlled synthesis of riboflavin in plants. Planta 226:147–58.

47. Waters BM, Amundsen K, Graef G. 2018. Gene expression profiling of iron deficiency chlorosis sensitive and tolerant soybean indicates key roles for phenylpropanoids under alkalinity stress. Frontiers in Plant Science 9:10.

48. Zamioudis C, Hanson J, Pieterse CM. 2014. β-Glucosidase BGLU42 is a MYB72-dependent key regulator of rhizobacteria-induced systemic resistance and modulates iron deficiency responses in Arabidopsis roots. New Phytologist 204: 368–79.

49. Ziegler J, Schmidt S, Strehmel N, Scheel D, Abel S. 2017. Arabidopsis transporter ABCG37/PDR9 contributes primarily highly oxygenated coumarins to root exudation. Scientific Reports 7: 3704.

50. Zocchi G, De Nisi P, Dell’Orto M, Espen L, Gallina PM. 2007. Iron deficiency differently affects metabolic responses in soybean roots. Journal of Experimental Botany 58: 993–1000.

